# KLF4 coordinates corneal epithelial apicobasal polarity and plane of cell division, and is downregulated in ocular surface squamous neoplasia

**DOI:** 10.1101/2020.01.08.899385

**Authors:** Anil Tiwari, Sudha Swamynathan, Vishal Jhanji, Shivalingappa K. Swamynathan

## Abstract

Apicobasal polarity (ABP) is an important feature of many epithelial cells, including those in the stratified squamous corneal epithelium (CE). Previously, we demonstrated that KLF4 promotes CE homeostasis by suppressing epithelial-mesenchymal transition (EMT) and TGF-β signaling. As dysregulation of TGF-β signaling affects ABP, we investigated the role of KLF4 in regulating cell polarity and plane of division by spatiotemporally regulated ablation of *Klf4* in adult ternary transgenic *Klf4^Δ/ΔCE^* (*Klf4*^*LoxP*/*LoxP*^/*Krt12*^*rtTA*/*rtTA*^/*Tet-O-Cre*) mouse CE. *Klf4*^*Δ*/*ΔCE*^ cells displayed decreased expression and mis-localization of apical polarity markers Pals1 and Crumbs1, apicolateral Par3, and basolateral Scribble. Cdc42 was upregulated, while Rac and Rho were mis-localized in the *Klf4*^*Δ*/*ΔCE*^ cytoplasm unlike their cortical expression in the control. Phalloidin staining revealed disrupted actin cytoskeleton in the *Klf4*^*Δ*/*ΔCE*^ cells. Survivin and phospho-histone-H3 immunostaining revealed a tilt in the plane of cell division favoring symmetrical divisions with vertical axis in the *Klf4*^*Δ*/*ΔCE*^ compared with predominantly asymmetrical divisions with horizontal axis in the control. Human ocular surface squamous neoplasia (OSSN) tissues displayed signs of EMT and loss of ABP markers PAR3, PALS1 and SCRIB, coupled with downregulation of KLF4. By demonstrating that *Klf4* ablation affects CE expression of ABP markers and Rho family GTPases, cytoskeletal actin organization and the plane of cell division, and that KLF4 is downregulated in OSSN tissues that display EMT and lack ABP, these results elucidate the key integrative role of KLF4 in coordinating CE cell polarity and plane of division, loss of which results in OSSN.

## Introduction

The corneal epithelium (CE), the anterior-most part of the eye that provides the transparent barrier function, is a self-renewing stratified squamous tissue comprised of basal proliferating cells that serve as a source for the differentiating suprabasal and terminally differentiated superficial cells which are eventually sloughed off. Cellular polarity-polarized distribution of protein complexes involved in cell-cell and cell-matrix interactions- is a fundamental determinant of epithelial cell properties (1–3). Epithelial cell polarity manifests as apicobasal polarity (ABP; polarization along the apicobasal axis that facilitates apical barrier formation and basal adhesion to basement membrane), and planar cell polarity (PCP; polarization along the orthogonal axis within the plane of the epithelium). Establishment and maintenance of correct cellular polarity is essential for all epithelial cells including the CE for their specialized cellular functions and homeostasis (2). A proper balance between CE cell proliferation and differentiation is essential for its homeostasis (4). Disruption of this balance results in severe visual impairments including epithelial erosion, dry eye, corneal fibrosis, and rare ocular tumors such as ocular surface squamous neoplasia (OSSN) (5). Though the role of PCP in CE cell migration has been investigated (6), little is known about the functions of ABP in CE. While studies in other stratified tissues like the skin suggest that ABP regulates the vectoral distribution of information from basal to apical cells (1), the molecular events that coordinate directionality and transmission of such information in the CE cells are poorly understood.

Epithelial ABP is established by the differential localization of three major membrane-associated protein complexes: the apical Crumbs complex comprised of Crumbs1 (Crb), Pals1 and PatJ, apicolateral Par complex consisting of Par6, Par3 and atypical PKC (aPKC), and the basolateral Scribble complex composed of Scribble, Dlg and Lgl (2,7). Components of Par complex interact with other apical polarity regulating factors such as Pals1 (8), which acts as a scaffold between PatJ and Crb to form a Crb/Pals1/PatJ complex localized at the tight junctions (9). ABP complex components also interact with the small GTPases Rho/Rac/Cdc42 that in turn help maintain F-actin cytoskeleton, providing structural stability to epithelial tissues (7,10,11). Besides maintaining ABP and cytoskeletal architecture, cell polarity proteins also facilitate directional spindle assembly and asymmetric cell divisions in stem and transiently amplifying cells which result in a daughter cell that undergoes differentiation while the other retains the renewal potential (12,13). Though it is widely recognized that the asymmetric localization of multiprotein complexes that demarcate the apical, lateral and basal aspects of an epithelial cell is evolutionarily conserved, how their expression is coordinated remains relatively understudied.

Previous studies from our lab and others identified Krüppel-like factor-4 (KLF4), one of the most abundantly expressed transcription factors in the CE, as a major determinant of CE properties (14,15). Corneal Klf4-target genes collectively promote CE structural stability and barrier functions while suppressing epithelial-mesenchymal transition (EMT) and TGF-β signaling (14,16–21). Though Klf4 is known to influence the intestinal epithelial cell ABP (22), its involvement in regulating the core ABP determinants is unexplored in stratified tissues like the CE. Given that (i) Klf4 is abundantly expressed in the CE where it upregulates the expression of tight and adherence junction components that play a key role in ABP (19), (ii) *Klf4* ablation results in EMT and increased TGF-β signaling commonly associated with compromised ABP and epithelial tumors (20,21), (iii) TGF-β-induced EMT is invariably associated with a loss of ABP (23), and (iv) decreased expression or mutations in *Klf4* are commonly associated with tumors (24,25) that display loss of core polarity components and altered plane of cell division (26), we predicted that Klf4 contributes to CE homeostasis by coordinating CE cell ABP and plane of division. Data presented in this report reveal that spatiotemporally regulated ablation of *Klf4* in the adult mouse CE affects (*i*) the expression and sub-cellular localization of core ABP markers Pals1, Crumbs1, Par3 and Scribble, (*ii*) expression and/or localization of Rho family GTPases, (*iii*) cytoskeletal F-actin organization, and (*iv*) the plane of cell division, elucidating the key integrative role of Klf4 in coordinating CE cell ABP and plane of division. Moreover, *KLF4* was downregulated in human OSSN tissues that displayed signs of EMT and loss of ABP, suggesting that mutations or altered expression of *KLF4* is a potential causative factor for OSSN.

## Results

### Apicobasal polarity is disrupted in *Klf4*^*Δ/ΔCE*^ CE

As reported previously (14,17,19,20), CE-specific ablation of *Klf4* resulted in hyperplasia concurrent with downregulation of CE markers keratin-12 (Krt12), tight junction protein-1 (Tjp1), and E-cadherin, reminiscent of EMT (Supplemental Fig. 1). Given that (i) TJP1 co-localizes with Par3 (27), (ii) EMT is associated with loss of epithelial ABP, and (iii) loss of adhesion molecules like E-cadherin and TJP1 impairs CE barrier function which in turn is associated with altered localization of Par3 complex (16,27), we hypothesized that the ablation of *Klf4* disrupts CE ABP. Consistent with that prediction, RT-qPCR revealed significant downregulation of *Par3*, *Pals1*, *Crumbs* and *Scrib* transcripts in *Klf4*^*Δ*/*ΔCE*^ compared with the control CE (Fig. 1 A). Corresponding decrease in Par3, Pals1 and Scrib protein expression in the *Klf4*^*Δ*/*ΔCE*^ corneas was confirmed by immunoblots (Fig. 1B). Immunofluorescent staining in the control CE revealed apicolateral cortical localization of Par3 and Pals1, apical localization of Crumbs and basolateral expression of Scrib indicating proper apical-basal polarization (Fig. 1). In contrast, *Klf4*^*Δ*/*ΔCE*^ corneas displayed sharply decreased expression of Par3, Pals1, Crumbs1 and Scrib (Fig. 1C). Collectively, these results suggest that the CE-specific ablation of *Klf4* results in downregulation of ABP markers and that Klf4 regulates the CE expression of a functionally related subset of proteins that play an important role in establishing and maintaining ABP.

**Fig. 1.**
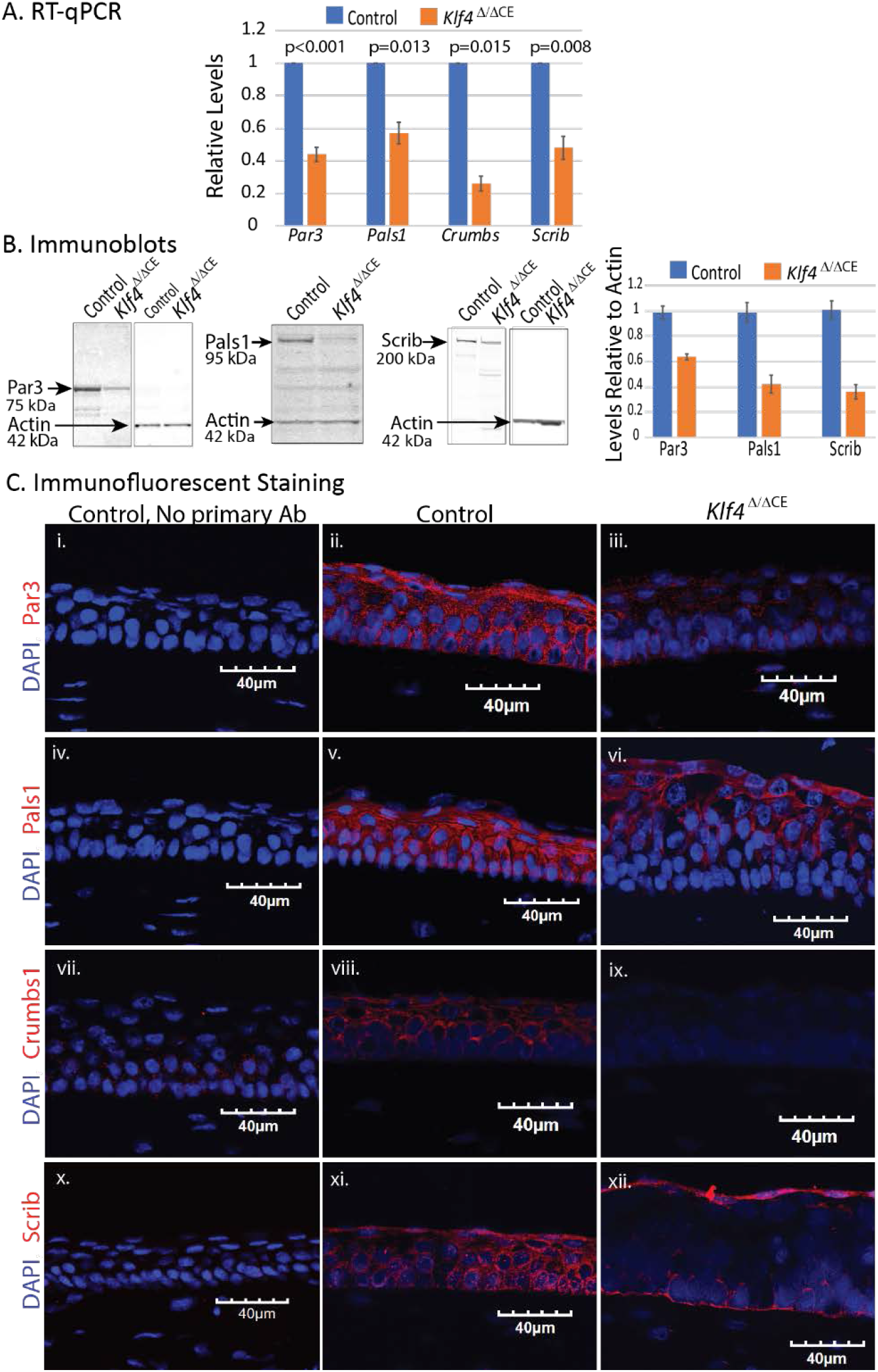
Apicobasal polarity is disrupted in *Klf4*^*Δ*/*ΔCE*^ corneal epithelium. (A) RT-qPCR reveals decreased expression of Par3 (n=4; p= 0.00058), Pals1 (n=4; p= 0.0134), Crumbs (n=4; p= 0.015) and Scrib (n=4; p=0.0081) in *Klf4*^*Δ*/*ΔCE*^ compared with the control corneas. (B) Immunoblots show decreased expression of Par3 (n= 4; p= 0.039), Pals1 (n= 4, p= 0.034) and Scrib (n=4; p=0.003) in *Klf4*^*Δ*/*ΔCE*^ corneas compared with the control. For densitometric quantitation, actin staining intensity was used as a loading control. All data are presented as mean ± ½ SEM. (C) Immunofluorescent stain for Par3 (i-iii), Pals1 (iv-vi), Crumbs (vii-ix) and Scrib (x-xii) in the *Klf4*^*Δ*/*ΔCE*^ corneas compared with the control. Corresponding no primary antibody controls are shown for each antibody (n=4; representative images shown).

### Rho GTPase localization and cytoskeletal actin organization is disrupted in the *Klf4^Δ/ΔCE^* CE

An array of signaling pathways participates in the regulation of ABP (28). Among them, Rho GTPase pathway is prominent, being implicated in diverse events that depend on cellular polarity. The best characterized members of the Rho family- Cdc42, Rho and Rac- play crucial roles in maintaining cellular structure and function by regulating the actin cytoskeletal organization (10,11,29,30). Therefore, we examined if Rho GTPase expression and/or subcellular localization is affected in *Klf4*^*Δ*/*ΔCE*^ CE concomitant with its disrupted ABP. We observed significantly increased levels of Cdc42 mRNA and protein in *Klf4*^*Δ*/*ΔCE*^ corneas compared with the control (Fig 2 A-B). Immunofluorescent stain further revealed strong cortical positioning of Cdc42 in both the control and the *Klf4*^*Δ*/*ΔCE*^ CE, with much more abundant expression in the *Klf4*^*Δ*/*ΔCE*^ cytoplasm (Fig 2C). Though RhoA and RhoB transcripts were moderately upregulated in the *Klf4*^*Δ*/*ΔCE*^ corneas (Fig. 2A), a commensurate increase in the protein levels was not observed by immunoblot using antibody against RhoA/B/C in *Klf4*^*Δ*/*ΔCE*^ corneas (Fig 2B). Immunofluorescent stain revealed diffuse cytoplasmic expression of RhoA/B/C and Rac1 in the *Klf4*^*Δ*/*ΔCE*^ compared with peripheral membrane-associated expression in the control CE (Fig 2C). Phalloidin stain revealed thick cortically localized F-actin cytoskeletal bundles in the control CE compared with those that were thin, lacked cortical localization and diffusely distributed in the *Klf4*^*Δ*/*ΔCE*^ CE cytoplasm (Fig 2C). Collectively, these results suggest that the loss of ABP in *Klf4*^*Δ*/*ΔCE*^ CE cells is accompanied by over-expression of Cdc42, loss of cortical localization of Cdc42, Rho and Rac1 and disruption of F-actin cytoskeletal network.

**Fig. 2.**
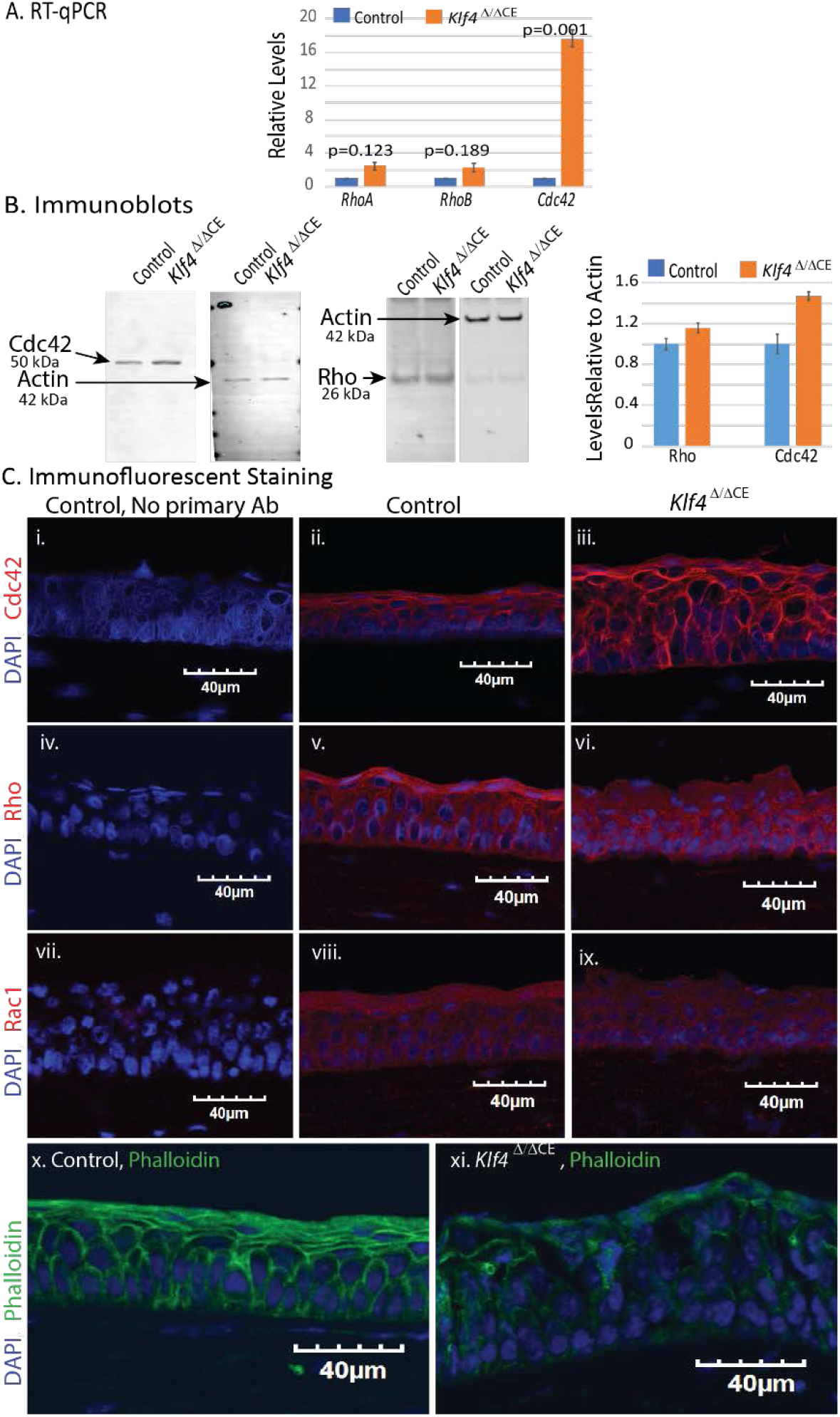
*Klf4*^*Δ*/*ΔCE*^ cells display increased expression and/or altered localization of Rho GTPases Cdc42, Rho and Rac. (A) RT-qPCR revealed a modest increase in *RhoA* and *RhoB* transcripts in the *Klf4*^*Δ*/*ΔCE*^ compared with the control corneas (*RhoA*: n=4; p=0.123 and *RhoB*; n=4; p=0.189) and a significant increase inCdc42 (n=4; p=0.001). (B) Immunoblots reveal increased expression of Cdc42 (n= 4; p= 0.016) in the *Klf4*^*Δ*/*ΔCE*^ corneas compared with the control. A similar increase was not observed in the Rho protein level (n=4; p=0.495). For densitometric quantitation, actin staining intensity was used as a loading control. All data are presented as mean ± ½ SEM. (C) Immunofluorescent stain shows increased expression of Cdc42 (i-iii) in the *Klf4*^*Δ*/*ΔCE*^ compared with the control CE. Rho (iv-vi) is predominantly localized to the control CE cell membranes, while it is also present in the cytoplasm and the nucleus in the *Klf4*^*Δ*/*ΔCE*^ CE. Rac1 (vii-ix) is membrane-associated in the control but is diffusely localized in *Klf4*^*Δ*/*ΔCE*^ CE cytoplasm. Staining with fluorescently tagged phalloidin (x-xi) revealed thick cortical F-actin bundles in control CE cells which were missing in the *Klf4*^*Δ*/*ΔCE*^ cells. (n= 4; Representative images shown).

### Klf4 promotes asymmetric, horizontal plane of division in basal CE cells

During cell division, mitotic spindle orientation and the plane of cell division are influenced by the cell’s polarity, which in turn is regulated by the asymmetric enrichment of ABP complex proteins (31). Given (*i*) the requirement for a proper balance between symmetric and asymmetric cell divisions during CE development and homeostasis (32), (*ii*) the influence of cellular polarity on plane of division (33), (*iii*) the dependence of stem cells on asymmetric cell division (34), and (*iv*) the role of Klf4 in maintenance of CE cell ABP described above, we next investigated if Klf4 is involved in regulating the CE plane of cell division. We evaluated the plane of cell division by immunofluorescent staining using anti-phospho-histone H3 (PH3) and anti-survivin antibodies (Fig. 3). A large fraction of the dividing cells in the control CE underwent asymmetrical division with a horizontal plane of division within 0-22.5° of the basement membrane (44% and 38%, relative to 23% and 26% symmetric divisions based on PH3 and survivin, respectively). Such events would presumably result in daughter cells with two different cell fates, i.e., proliferation and differentiation, essential for stratification. (Fig. 3) (35). In contrast, *Klf4*^*Δ*/*ΔCE*^ CE displayed a tilt in the plane of cell division favoring symmetrical divisions with a vertical axis within 67.5-90° of the basement membrane (39% and 47% relative to 16% and 21% asymmetric divisions in the control CE based on PH3 and survivin, respectively), that is expected to produce the relatively increased number of dividing cells within *Klf4*^*Δ*/*ΔCE*^ CE as reported earlier (Fig. 3) (17). No significant difference was observed in the number of cells undergoing oblique plane of division within 22.5-67.5° of the basement membrane in WT (33% and 36% based on PH3 and survivin staining) and *Klf4*^*Δ*/*ΔCE*^ CE (45% and 32% with PH3 and survivin staining, respectively).

**Fig. 3.**
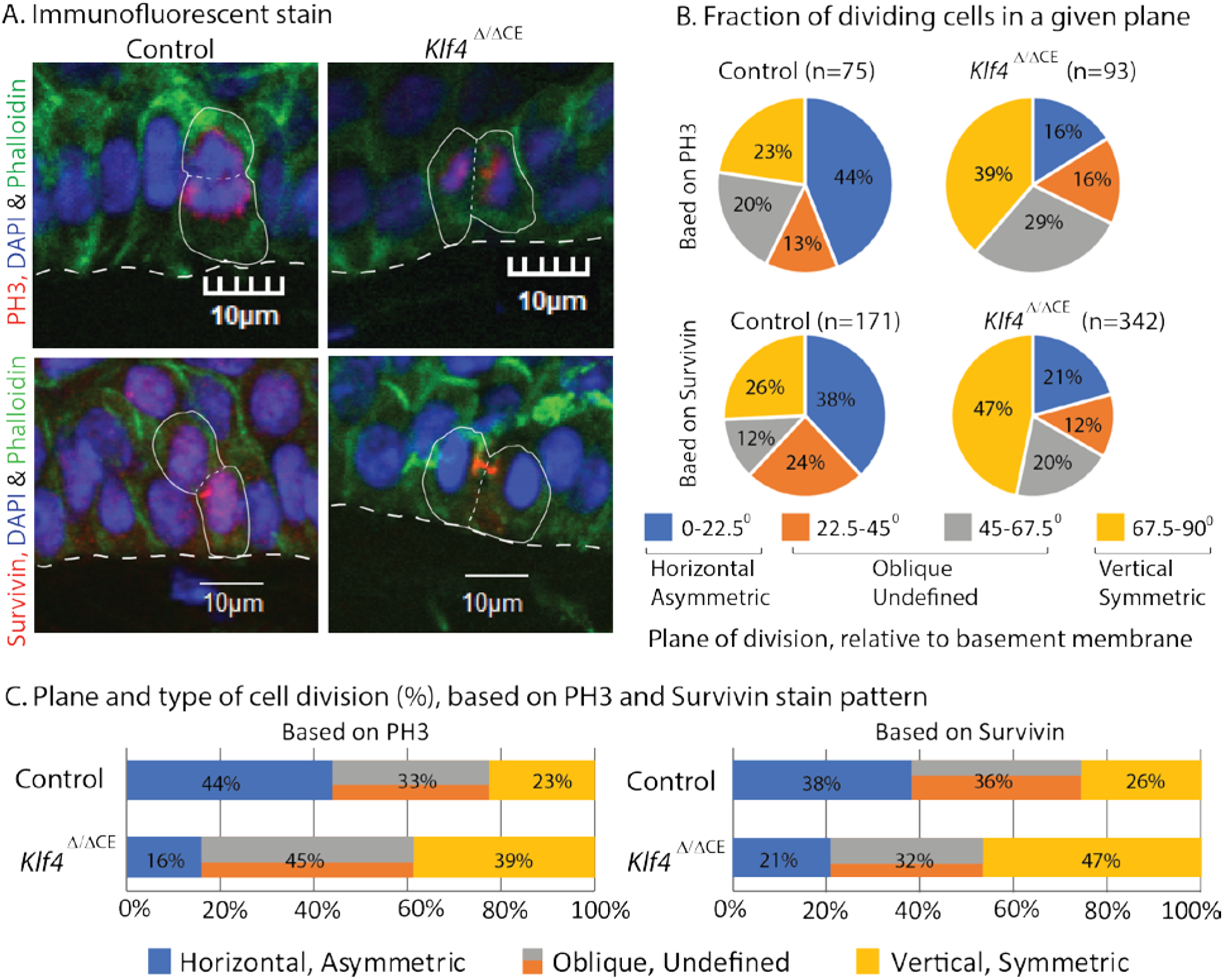
*Klf4*^*Δ*/*ΔCE*^ CE cells favor vertical plane of division, unlike horizontal plane of division in the control CE. (A) Representative images of immunofluorescent stain with anti-phospho-histone-H3 (PH3) and anti-survivin antibodies in the control and *Klf4*^*Δ*/*ΔCE*^ corneas. Basal CE cells stained for PH3 or Survivin (Red), F-actin (Phalloidin; green) and nuclei (DAPI; blue) are shown with the plane of division marked by a thin dotted line relative to the basement membrane (dotted white line). (B and C) Distribution of the plane of division in control and *Klf4*^*Δ*/*ΔCE*^ CE quantified by analyzing 4 adjacent images from the central CE in 4 sections each from 5 different eyeballs. (B) Pie charts displaying the distribution of the angle of plane of division relative to basement membrane in control and *Klf4*^*Δ*/*ΔCE*^ CE. The number of nuclei counted in each condition (n value) and the percentage of cells falling in each group is indicated. (C) Corresponding histograms showing the relative decrease in asymmetric division (0-22.5º) and increase of symmetric division (67.5-90º) in *Klf4*^*Δ*/*ΔCE*^ CE compared with the control. Planes of division in the 22.5-67.5º range which have roughly equal potential to result in symmetric or asymmetric divisions were considered oblique.

### EMT and loss of ABP in human ocular surface squamous neoplasia (OSSN) is associated with downregulation of KLF4

Previously, we reported that CE-specific ablation of *Klf4* results in defects that resemble OSSN (17,20,21). To determine if OSSN is indeed accompanied by EMT, we obtained surgically excised human tissues suspected of OSSN, and ascertained OSSN by histology (Fig. 4A). RT-qPCR revealed significant downregulation of *KRT12*, a CE-specific marker and a KLF4-target gene (14,16), coupled with upregulation of EMT-inducers *TGF-β1* and *TGF-β2*, and EMT transcription factors *SLUG*, *ZEB1*, *ZEB2*, and *TWIST1* in OSSN compared with the normal CE (Fig. 4B, Supplementary Table S1). Immunofluorescent stain confirmed abundant expression of KRT12 in the normal CE, which was sharply decreased in OSSN tissues (Fig. 4C, Supplementary Table S1). Immunostaining also revealed an abnormally high frequency of Ki67+ and survivin+ cells in OSSN tissue compared with the normal control, consistent with the high rate of proliferation in OSSN tissues (Fig. 4C, Supplementary Table S1). Next, we confirmed the loss of epithelial features in OSSN tissues by evaluating the expression and localization of E-cadherin and β-catenin. Immunostaining revealed that both E-cadherin and β-catenin are abundantly expressed and properly localized to the cell membranes in the control CE where they form a part of the adherens junctions (Fig. 5). In contrast, E-cadherin was diffusely localized in the cytoplasm and β-catenin was sharply downregulated and abnormally localized in nuclei, consistent with EMT in OSSN samples (Fig. 5, Supplementary Table S1).

**Fig. 4.**
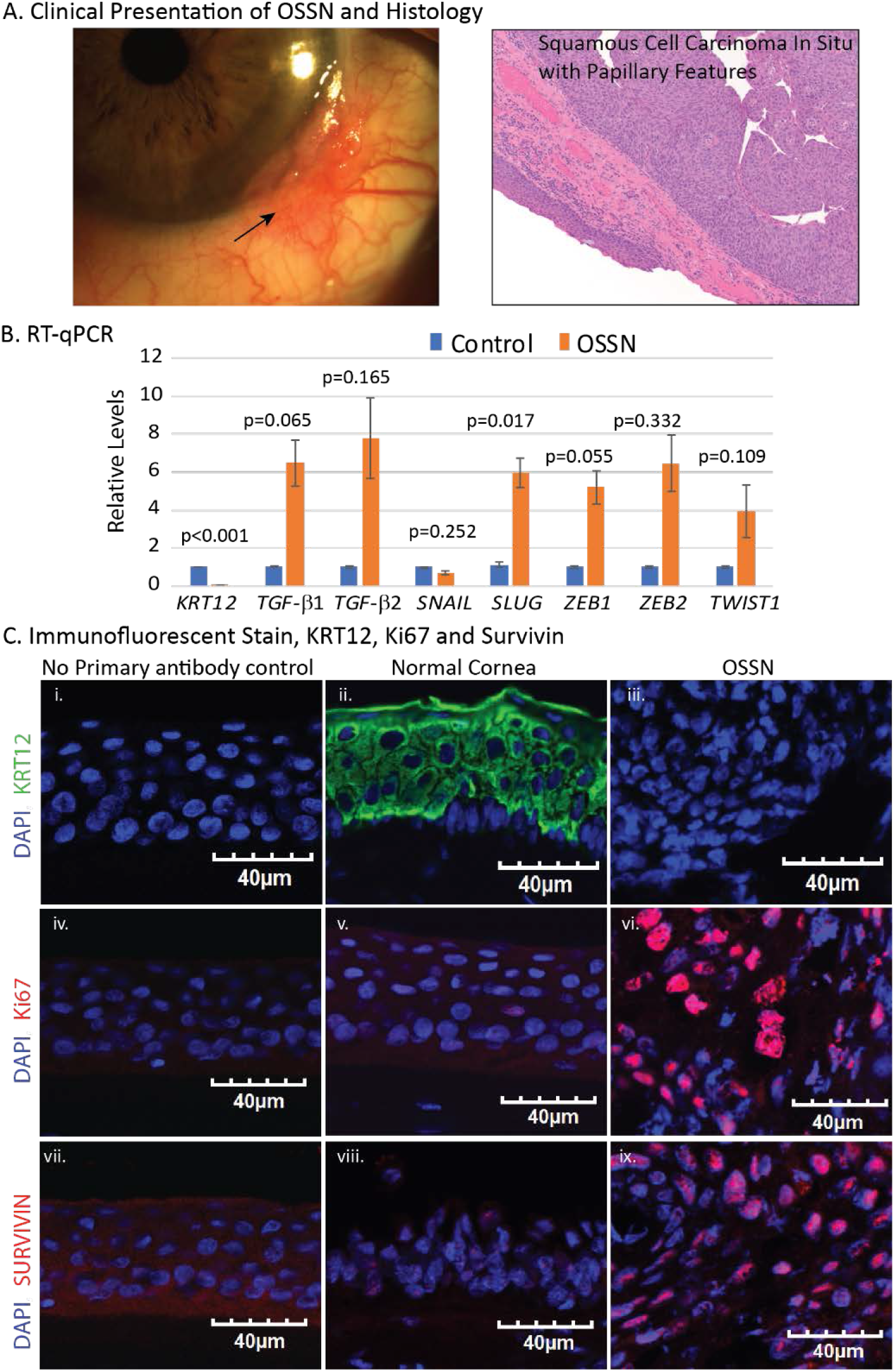
Signs of EMT in OSSN tissue. A. En face image of OSSN and histology of excised tissue. Histology suggests squamous cell carcinoma in situ with papillary features. B. RT-qPCR reveals downregulation of CE-specific marker KRT12 and upregulation of EMT-inducers TGF-β1 and TGF-β2, as well as EMT-transcription factors SLUG, ZEB1, ZEB2, and TWIST1 (n=4; p values shown). C. Immunofluorescent stain reveals (i-iii) abundant expression of KRT12 (green) in (ii) normal CE but not (iii) OSSN, (iv-ix) abnormally high number of Ki67+ and Survivin+ cells in (red; vi and ix) OSSN relative to far fewer Ki67+ and Survivin+ cells in normal CE (red; v and viii). No primary antibody control for each antibody used is shown (n= 3; Representative images shown).

**Fig. 5.**
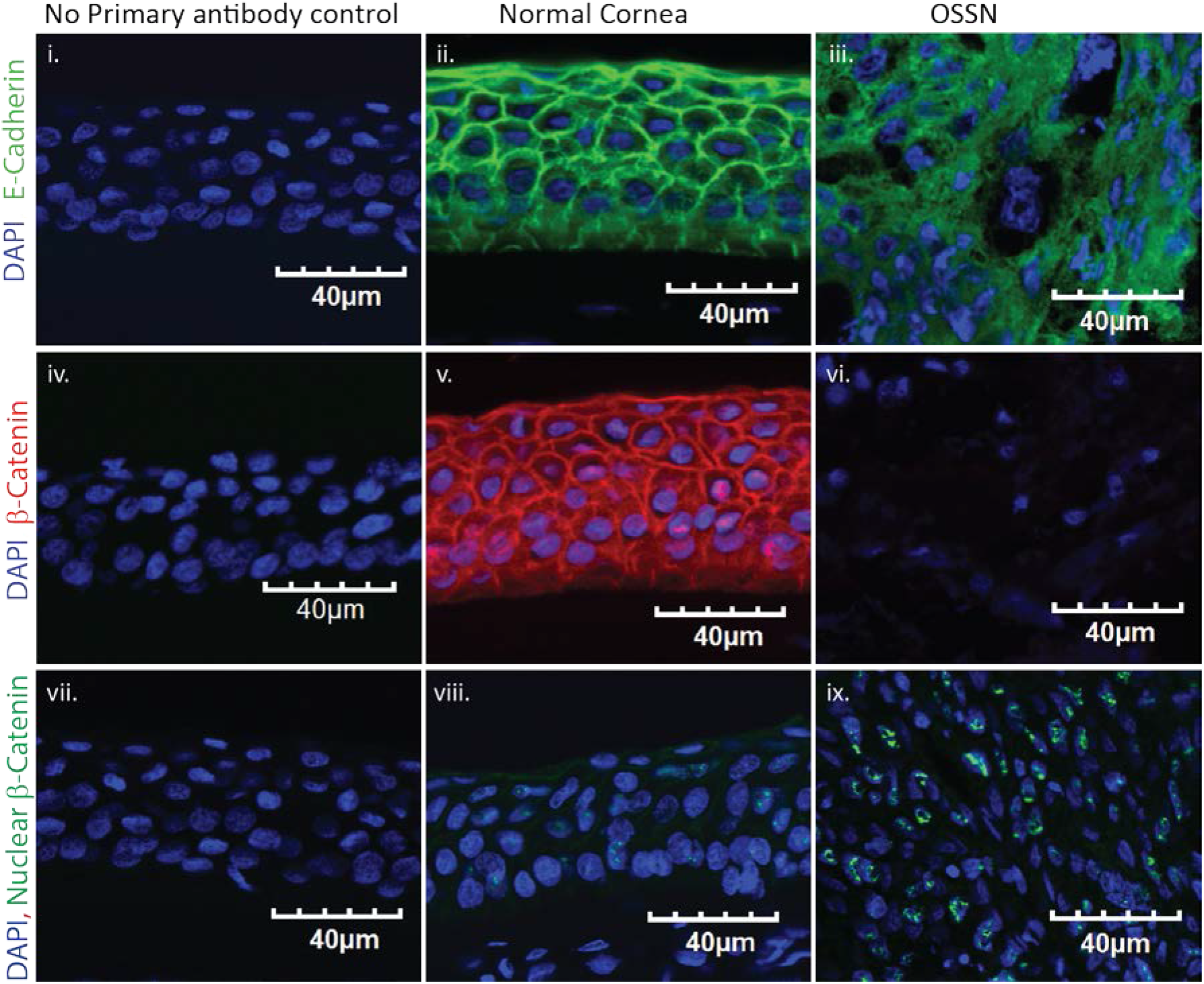
Loss of epithelial properties in OSSN. Immunofluorescent stain reveals abundant expression and proper membrane localization of (ii) E-Cadherin and (v) β-Catenin in the normal CE, compared with sharply decreased expression of (iii) E-Cadherin that is diffusely localized in the cytoplasm, and (vi) β-Catenin in OSSN. Immunostaining with an antibody that specifically detects the nuclear β-Catenin (green) revealed (ix) strong nuclear presence of β-Catenin in many OSSN cells compared with a (viii) a faint expression in far fewer cells in the normal CE nuclei. No primary antibody control for each antibody used is shown (n= 3; Representative images shown).

Next, we determined if EMT in OSSN is also associated with a loss of ABP by testing the expression and localization of PAR3, PALS1 and SCRIB. RT-qPCR revealed significant downregulation of *PAR3*, *PALS1* and *SCRIB* in OSSN compared with the control CE (Fig. 6A, Supplementary Table S1). Consistently, immunostaining revealed that PAR3, PALS1 and SCRIB are sharply downregulated and diffusely localized in the cytoplasm of OSSN cells, unlike normal apicolateral localization of PAR3 and PALS1, and basolateral expression of SCRIB in the control CE (Fig. 6B). Collectively, these data reveal that OSSN cells display different signs of EMT including elevated cell proliferation, downregulation and disrupted localization of epithelial markers, and loss of ABP determinants. Considering that each of these features was also observed in the *Klf4*^*Δ*/*ΔCE*^ corneas with CE-specific ablation of *Klf4*, we hypothesized that EMT and loss of ABP in OSSN is an outcome of downregulation of KLF4. Consistent with this prediction, RT-qPCR revealed significant downregulation of KLF4 in OSSN tissues, which was confirmed by immunofluorescent stain (Fig. 7, Supplementary Table S1). Collectively, these results demonstrate that KLF4 upregulates the expression of ABP complex components thereby promoting asymmetric pattern of division required for CE stratification and homeostasis, and that KLF4 is downregulated in OSSN tissues which display signs of EMT and loss of ABP.

**Fig. 6.**
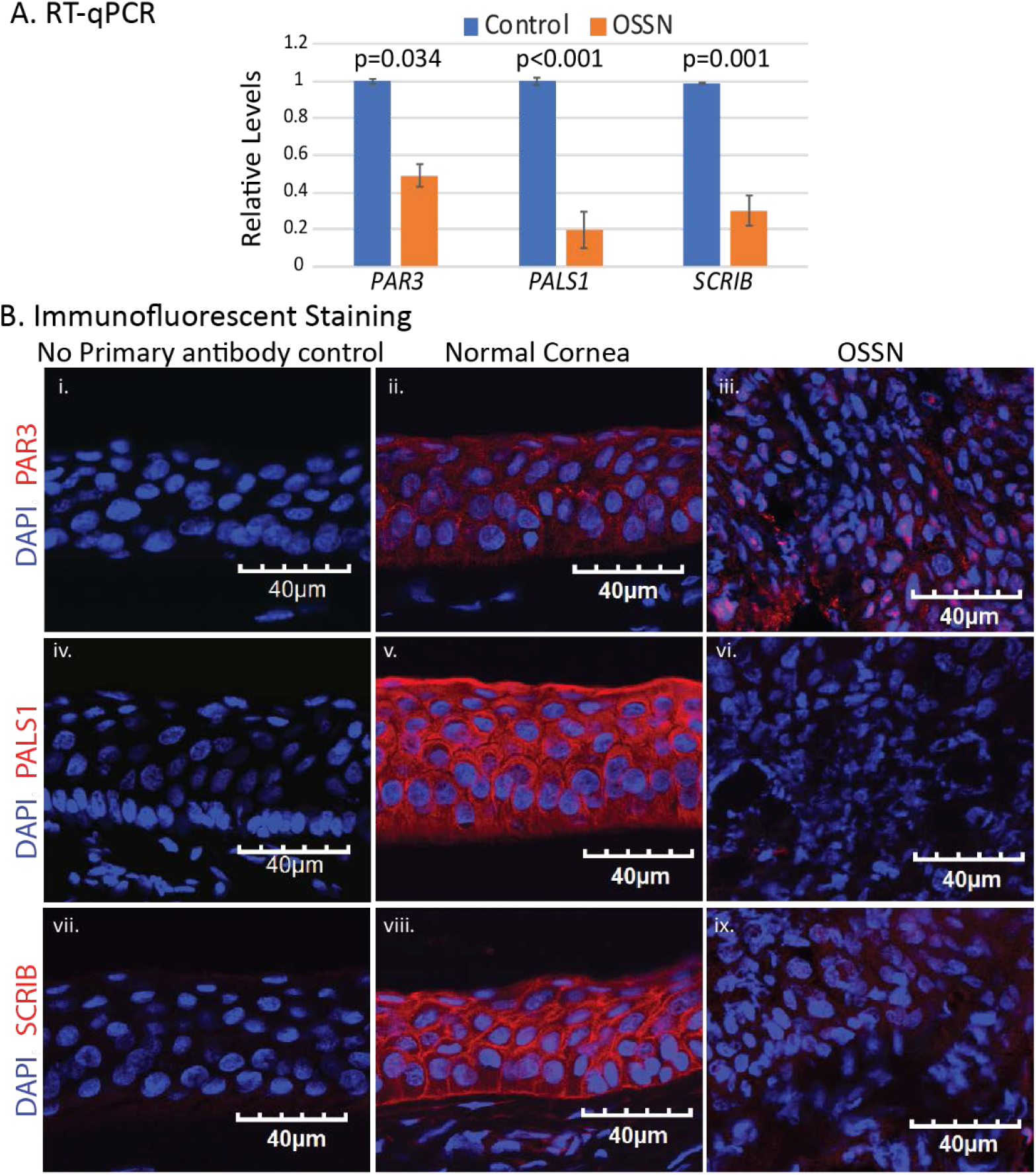
Loss of ABP in OSSN. A. RT-qPCR reveals significant downregulation of *PAR3*, *PALS1* and *SCRIB* transcripts in OSSN compared with the normal CE. N=4; p values shown on the graph. B. Immunofluorescent staining reveals that the normal CE displays abundant expression and proper localization of (ii) PAR3, (v) PALS1, and (viii) SCRIB, that is sharply downregulated and mis-localized in the OSSN tissue (iii, vi and ix, respectively). No primary antibody control for each antibody used is shown (n= 3; Representative images shown).

**Fig. 7.**
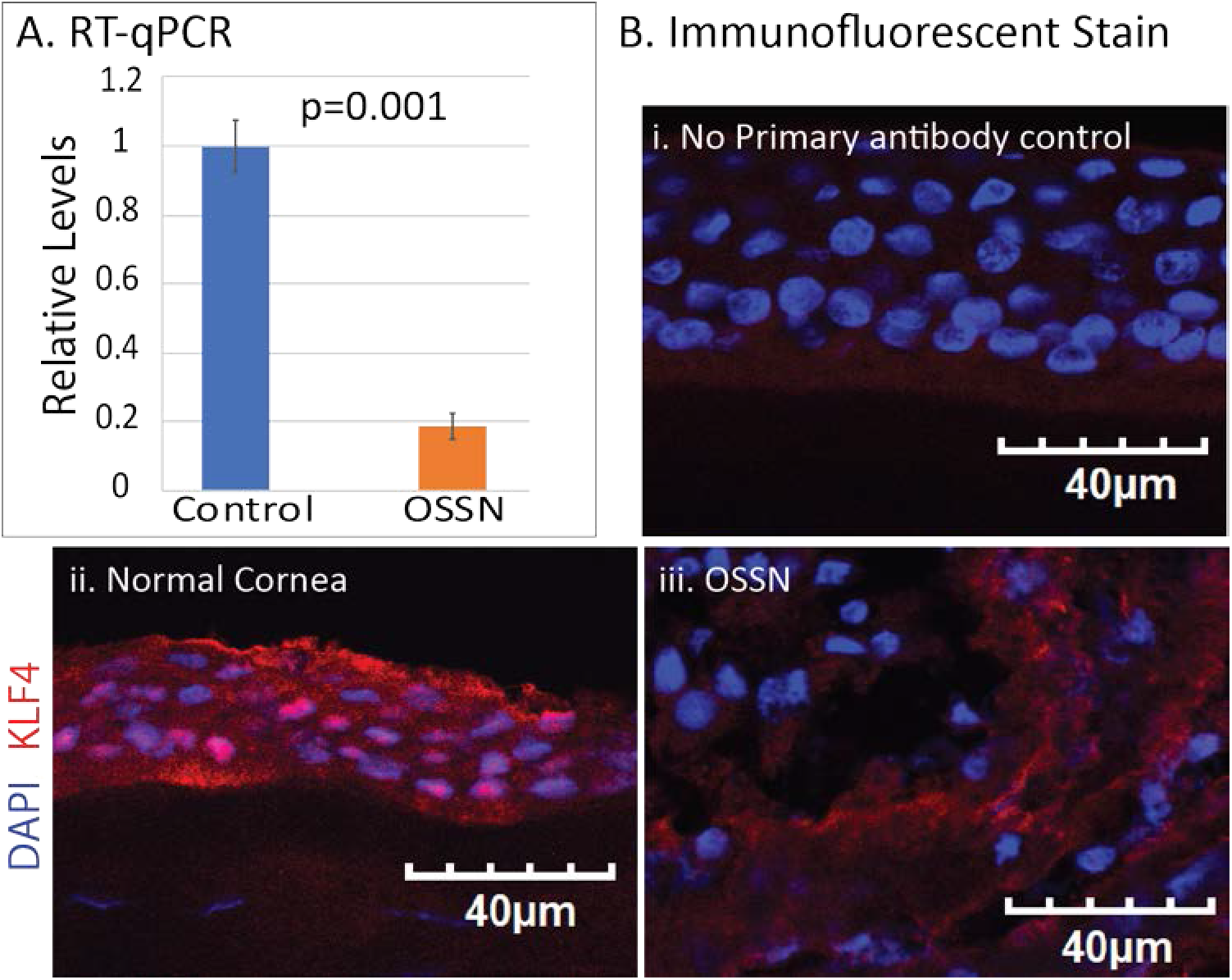
KLF4 is downregulated in OSSN. A. RT-qPCR reveals significant downregulation of *KLF4* transcripts in OSSN compared with the normal CE (n=4; p values shown on the graph). B. Immunofluorescent staining reveals that the normal CE displays abundant expression and proper nuclear localization of KLF4 (panel ii) that is sharply downregulated and mis-localized in the OSSN tissue (panel iii). No primary antibody control is shown (panel i) (n= 3; Representative images shown).

## Discussion

Previously, we reported that CE-specific ablation of *Klf4* results in (i) EMT coupled with loss of CE barrier function and hyperplasia (14,17,20), and (ii) activation of canonical TGF-β signaling and downregulation of cell cycle inhibitors favoring increased proliferation (21). The data presented in this report elucidate the crucial role of Klf4 in orchestrating the CE stratification and homeostasis by coordinating the expression and localization of ABP determinants, and plane of CE cell division (Fig. 8). Our data also demonstrate that (*i*) human OSSN tissues display signs of active EMT manifested as increased expression of TGF-β and EMT transcription factors, (*ii*) EMT in OSSN tissues is concurrent with loss of ABP, and (*iii*) KLF4 and its target genes KRT12, E-cadherin and β-catenin are significantly downregulated in OSSN. Collectively, these results demonstrate that KLF4 plays a key integrative role in coordinating CE cell polarity and plane of cell division, and that the loss of this key function results in OSSN with potentially devastating consequences on sight (Fig. 8).

**Fig. 8.**
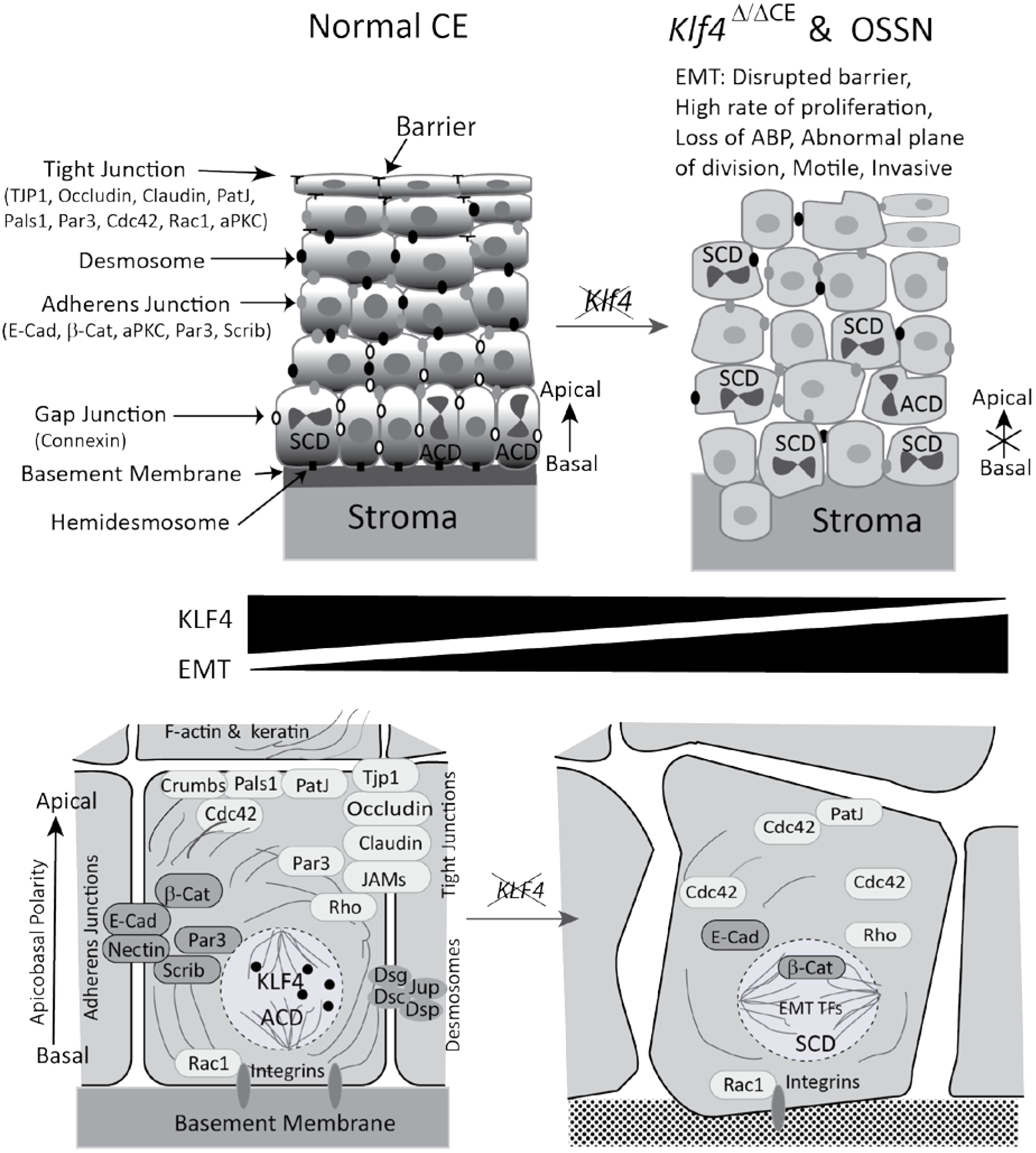
Schematic summarizing the findings described in this study. Mouse CE-specific ablation of *Klf4* resulted in disruption/downregulation of ABP core complex components and Rho family GTPases altering actin cytoskeletal organization and favoring the symmetric plane of cell division (SCD) compared with the asymmetric cell divisions (ACD) favored in the control. Sharply downregulated expression of ABP core complex components in the human OSSN samples was coupled with decreased expression of KLF4. These changes are shown in greater detail in the lower panels. By demonstrating that KLF4 promotes the expression of ABP complex components thereby wiring the asymmetric pattern of division essential for CE self-renewal and differentiation, these results highlight the importance of KLF4 in CE stratification and homeostasis.

Unlike simple epithelial cells where ABP is defined by the attachment of the cells to the basement membrane on the basal side, mechanisms that regulate the distribution of polarity-determining components across the stratified epithelial tissue are poorly understood (1). Initial formation of a polarized stratified epithelium, and its homeostatic maintenance in the later adult stage involves three crucial events: (*i*) establishment of ABP across different layers; (*ii*) formation of apical tight junctions and apicolateral adherence junctions; and (*iii*) proper positioning of the mitotic spindle to enable asymmetric cell division (36). CE-specific ablation of *Klf4* resulted in downregulation of tight and adherence junction proteins Tjp1 and E-cadherin, respectively, suggesting a role for Klf4 in regulating ABP (17,20). Though the role of Klf4 in regulating the expression of tight- and adherence-junction components is well defined (16,19), its involvement in regulating ABP determinants Crumbs, Pals1, Par3 and Scribble was not described previously. Our observation that the loss of Klf4 results in the loss of ABP provides the first-ever evidence that Klf4 plays a crucial role in maintaining the polarized nature of the stratified CE.

Though it is well established that different ABP complex components including Crumbs, Par and Scribble are direct transcriptional targets of EMT inducers such as TGF-β (23,37), and that the epithelial ABP is disrupted with the onset of EMT (38), the underlying mechanistic basis for this disruption was not known. The CE-specific ablation of *Klf4* results in suppression of epithelial genes, and induction of EMT facilitated by robust TGF-β-signaling (17,20,21). Although the downregulation of ABP complex components in the *Klf4*^*Δ*/*ΔCE*^ CE described here suggests a supportive role for Klf4 in maintaining ABP, whether this downregulation is a direct effect of the absence of Klf4, or an indirect manifestation of the EMT mediated by upregulation of TGF-β reported earlier (20,21), remains to be established.

Establishment and maintenance of cell polarity requires efficient crosstalk between a complex network of different signaling pathways including that involving Rho family GTPase proteins (10,39,40). Rho GTPases regulate cell shape and surface dynamics by orchestrating the communication between the cytoskeleton, contractile actin cortex, and the plasma membrane (40). Among Rho GTPases, the expression of Cdc42 which contributes to the apical polarity by interacting with Par3 (41), and regulates nucleation of actin filaments (42) was upregulated in *Klf4*^*Δ*/*ΔCE*^ corneas, consistent with the ability of Klf4 to inhibit Cdc42 expression via WNT5A (43). Though there was no appreciable change in Rho expression in *Klf4*^*Δ*/*ΔCE*^ CE, its localization was altered compared with the cortical organization in the control. Together, these results suggest that Klf4 coordinates ABP by regulating the expression and cortical localization of Rho GTPases, which promote nucleation and cortical arrangement of F-actin (11,30,40).

The data presented here demonstrate that CE-specific ablation of *Klf4* results in loss of ABP which is essential for epithelial stratification and homeostasis. Asymmetric distribution of polarity determinants also governs the mitotic spindle orientation and promotes the asymmetric pattern of cell division, creating a pool of proliferating and differentiating cells aiding in the process of self-renewal in stratified tissues (39,44). (35). The current data elucidate that similar to other stratified tissues such as the epidermis (35), the WT CE display more asymmetric divisions- a condition necessary for stratification. In contrast, the *Klf4*^*Δ*/*ΔCE*^ corneas displayed a tilt in the plane of division favoring vertical plane of division expected to result in symmetrical daughter cells that retain the potential to further proliferate. This results in excessive proliferation and compromised differentiation as observed in different tumors involving mutations and/or deletions of *KLF4* (25,45–47). Consistent with this scenario, the absence of Klf4 potentially drives the *Klf4*^*Δ*/*ΔCE*^ CE towards OSSN, a group of ocular surface tumors characterized by hyperproliferation and aggressive migration (48). Our previous findings that *Klf4* ablation results in EMT via elevated TGF-β signaling, coupled with the current data that *Klf4* ablation results in the loss of ABP favoring vertical plane of division in the basal CE cells that in turn creates a symmetrical pool of proliferating cells suggest that Klf4 contributes to CE stratification and homeostasis by promoting correct ABP and asymmetrical plane of division. Unabated EMT and loss of ABP in OSSN tissues concomitant with the significant downregulation of *KLF4* is consistent with this important role for KLF4.

In summary, our current findings highlight the important role of KLF4 in promoting CE stratification and homeostasis by regulating the proper expression of the ABP complex components and Rho GTPases thereby ensuring the asymmetrical pattern of cell division essential for CE self-renewal and differentiation. By demonstrating that *Klf4* ablation affects CE expression of ABP markers and Rho family GTPases, cytoskeletal actin organization and the plane of cell division, and that KLF4 is downregulated in OSSN tissues that lack ABP and display EMT, these results elucidate the key integrative role of KLF4 in coordinating CE cell polarity and plane of division, loss of which results in OSSN (Fig. 8).

## Experimental Procedures

### Animals

All experiments were performed in accordance with the University of Pittsburgh Institutional Animal Care and Use Committee (IACUC Protocol # 17019882 titled ‘Role of Krüppel-like factors in the ocular surface’, PI: Swamynathan) and the ARVO Statement on the Use of Animals in Ophthalmic and Vision Research. All studies were conducted with 8- to 10-week-old mice. Spatiotemporal ablation of *Klf4* in CE was achieved by feeding ternary transgenic *Klf4*^*Δ*/*ΔCE*^ (*Klf4*^*LoxP*/*LoxP*^/ *Krt12*^*rtTA*/*rtTA*^/ *Tet-O-Cre*) mice with doxycycline (Dox) chow for at least a month. Age- and sex-matched littermates fed with regular chow served as control groups.

### Collection and processing of human normal corneas and OSSN samples

Normal human corneas were sourced from donor corneal tissues rejected for transplants, following the procedures approved by the University of Pittsburgh Committee for Oversight of Research and Clinical Training Involving Decedents (CORID ID # 889; titled Krüppel-like factors in the corneal epithelium, PI: Swamynathan). Human OSSN samples were collected following the Institutional Review Board (IRB)-approved protocol (# PRO-18100052, titled Ocular Surface Squamous Neoplasia, PI: Jhanji).

### Total RNA isolation and RT-qPCR

Total RNA was isolated from dissected mouse corneas or OSSN tissues using EZ-10 spin column total RNA mini-prep kit (Bio Basic, Inc., Amherst, NY, USA). Isolated RNA (500ng) was used for cDNA synthesis with mouse Maloney leukemia virus reverse transcriptase (Promega, Madison, WI, USA). SYBR Green RT-qPCR gene expression assays were performed in triplicate in an ABI StepOne Plus thermocycler using appropriate endogenous controls (Applied Biosystems, Foster City, CA, USA). The sequence of oligonucleotide primers used for RT-qPCR is presented in Supplementary Table S2.

### Immunoblots

Antibodies used in study are listed in Supplementary Table S3. Dissected *Klf4*^*Δ*/*ΔCE*^ or control corneas were homogenized in urea buffer (8.0 M urea, 0.8 % Triton X-100, 0.2 % SDS, 3 % β-mercaptoethanol, and protease inhibitors) and clarified by centrifugation. 20 µg of total protein in the supernatant was separated on 4-12% gradient polyacrylamide gels using 3-(N-morpholino) propanesulfonic acid (MOPS)/2-(N-morpholino) ethanesulfonic acid (MES) buffer and blotted onto polyvinylidine fluoride (PVDF) membranes. The membranes were blocked for 1h at room temperature, incubated overnight at 4°C with appropriate dilution of primary antibody, washed thrice with phosphate buffered saline (PBS) containing 0.1% Tween-20 (PBST) for 5 min each, incubated with fluorescently labeled secondary antibody (goat anti-rabbit IgG, or donkey anti-goat IgG) for 1 h at 23°C, washed three times with PBST for 5 min each followed by a wash with PBS to remove traces of Tween-20. Blots were scanned on Odyssey scanner (Li-Cor Biosciences, Lincoln, NE, USA), and densitometric measurements of the immunoreactive band intensities performed using Image J software (http://imagej.nih.gov/ij/; provided in the public domain by the National Institutes of Health, Bethesda, MD, USA). β-Actin was used as a loading control for normalizing the data.

### Immunofluorescent staining

Eight-micrometer-thick sections from OCT-embedded OSSN tissues, *Klf4*^*Δ*/*ΔCE*^ or control eyeballs were fixed in buffered 4% paraformaldehyde for 10 min at 23°C, washed thrice for 5 min each with PBS (pH 7.4), permeabilized (0.1% Triton X-100 in PBS) when necessary followed by three washes of 5 min each with PBS, treated with glycine for 20 min, washed thrice with PBS, blocked (10% goat or donkey serum in PBS) for 1 h at 23°C in a humidified chamber, washed twice with PBS for 5 min each, incubated with the appropriate dilution of the primary antibody for 2 h at 23°C or overnight at 4°C, washed thrice with PBS for 5 min each, incubated with appropriate secondary antibody (Alexafluor 546-coupled goat anti-rabbit IgG, Alexafluor 488-coupled goat anti-mouse IgG or Alexafluor 488-coupled donkey anti-goat IgG; Molecular Probes, Carlsbad, CA, USA) at a 1:400 dilution for 1 h at 23°C, washed thrice with PBST, counterstained with 4,6- diamidino-2-phenylindole (DAPI), mounted with Aqua-Poly/Mount (Polysciences, Warrington, PA, USA), and imaged using Olympus IX81 microscope (Olympus America, Inc.).

### Analysis, measurement and quantification of spindle orientation

WT and *Klf4*^*Δ*/*ΔCE*^ CE sections were immunofluorescently stained with anti-survivin and anti-phospho-histone H3 (PH3) antibodies to identify the mitotic cells, counterstained with DAPI, and the immunostaining pattern was used to determine the plane of division in the basal epithelial cells. Cells were taken into consideration only if both the daughter nuclei surrounding the survivin/PH3 immunostaining could be clearly identified. Distribution of the plane of division was quantified by analyzing 4 adjacent images from the central CE in 4 sections each from 5 different control and *Klf4*^*Δ*/*ΔCE*^ eyeballs. Mean counts were obtained in a blinded fashion from 75 and 93 nuclei that stained positive for PH3, and 171 and 342 nuclei that stained positive for survivin, respectively, from the WT and *Klf4*^*Δ*/*ΔCE*^ CE. The angel of division was calculated by plotting a line passing through the centers of the two nuclei relative to the basement membrane. The angel of division is represented in 22.5° increments from 0°-90°. Cell division positioned at 0-22.5° relative to the basement membrane were considered horizontal (asymmetrical), and those at 67.5-90° as vertical (symmetrical), with the remaining oriented at 22.5-67.5° considered as oblique.

### Statistical Analysis

The results presented here are representative of at least three independent experiments and shown as mean ± standard error of mean (SEM). Statistical significance was tested by Student’s t-test, with p≤ 0.05 considered statistically significant.

## Acknowledgements

We thank Mr. John Gnalian for technical help, Ms. Kate Davoli (Tissue Culture and Histology Core Module) for help with histology, Ms. Kira Lathrop (Imaging Core Module) for help with imaging, Dr. Xiangyun Wei for the generous gift of Crumbs1 antibody, and Dr. Debasish Sinha for critical comments on the manuscript.

## Author Contributions

A.T. designed and conducted the experiments, interpreted the results and prepared the manuscript. S.S. conducted the experiments and interpreted the results. V.J. collected OSSN samples, provided clinical evaluation and histology of OSSN samples. S.K.S. conceived the study, designed the experiments, interpreted the results and prepared the manuscript. All authors read and edited the manuscript and agree with its publication in its present form.

## Footnotes

This work was supported by NIH grants: R01 EY026533 (SKS), NEI core grant P30 EY08098; by unrestricted grants from Research to Prevent Blindness and the Eye and Ear Foundation of Pittsburgh.

## The abbreviations used are

OSSN: ocular surface squamous neoplasia
CE: corneal epithelium
KLF4: Krüppel-like factor-4
EMT: epithelial-mesenchymal transition
ABP: apicobasal polarity
PCP: planar cell polarity
TGF-β: transforming growth factor-beta
PVDF: polyvinylidine fluoride
MOPS: 3-(N-morpholino) propanesulfonic acid
MES: 2-(N-morpholino) ethanesulfonic acid
DAPI: 4,6-diamidino-2-phenylindole
OCT: Optimal cutting temperature compound
SEM: standard error of mean

## Supplemental Data

**Supplemental Fig. 1.**
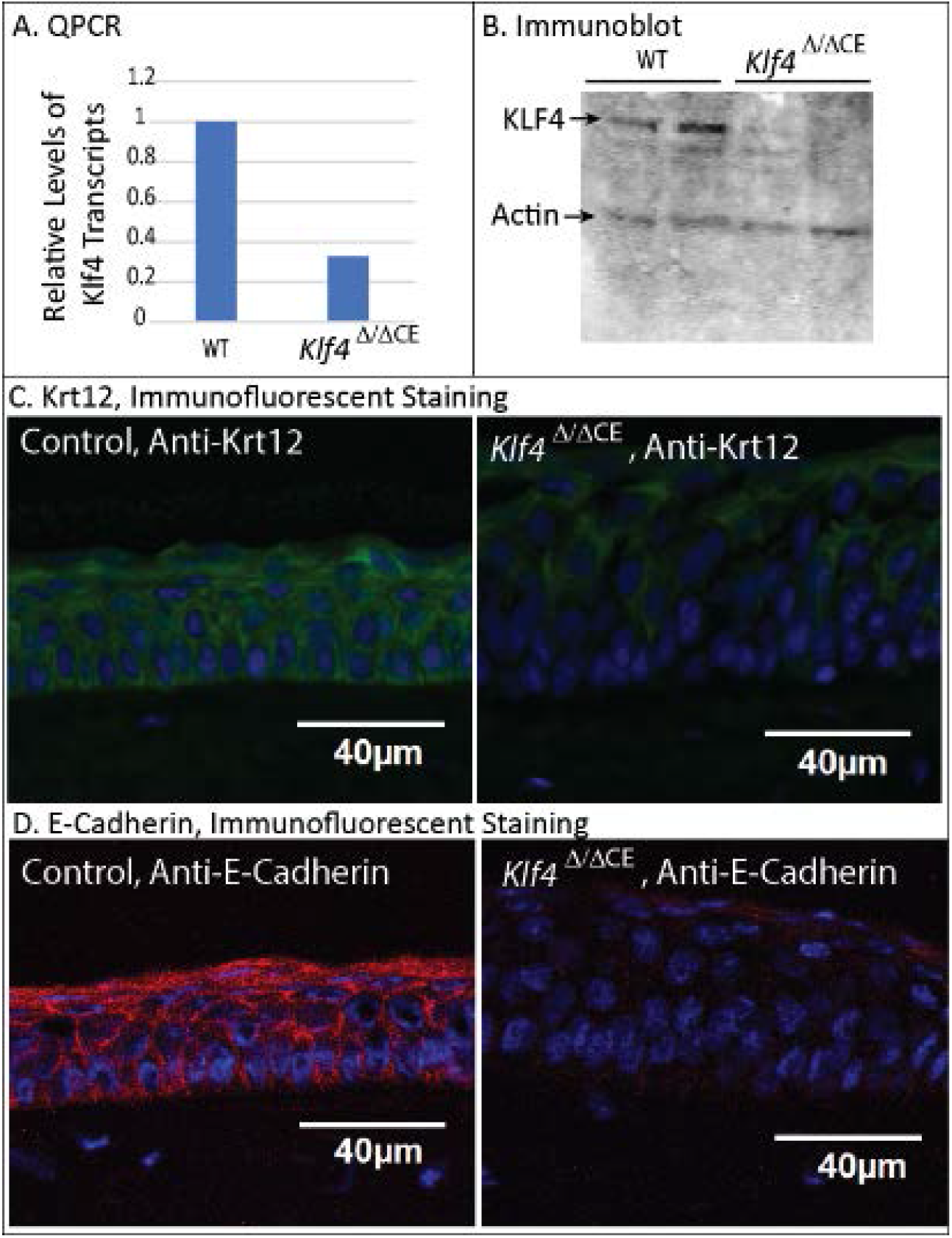
CE-specific ablation of *Klf4* results in loss of CE marker Krt12 and adherence junction protein E-Cadherin. (A) Confirmation of *Klf4* ablation by RT-qPCR (B) Confirmation of *Klf4* ablation by immunoblot. (C-D) Immunofluorescent stain showing downregulation of CE marker Krt12 (C) and adherence junction protein E-cadherin (D), both direct targets of Klf4, in *Klf4*^*Δ*/*ΔCE*^ CE (n=4; Representative images shown).

**Supplementary Table S1.** Clinical grade of severity, age and sex of OSSN patients, and relative expression levels of select genes in four different OSSN samples compared with the control tissues, determined by RT-qPCR and immunostaining. DR, Downregulated; UR, Upregulated; NT, Not tested.

| OSSN Sample # | 1 | 2 | 3 | 4 |
| --- | --- | --- | --- | --- |
| Severity Grade | Moderate | Mild | Severe | Mild |
| Age (Years) | 81 | 30 | 36 | 52 |
| Sex (M/F) | M | F | M | F |
| **Relative expression levels of select genes determined by RT-qPCR** |  |  |  |  |
| *KLF4* | 0.18 | 0.45 | 0.06 | 0.04 |
| *KRT12* | 0.17 | 0.01 | 0.14 | 0.003 |
| *SNAIL* | 1.5 | 0.2 | 0.17 | 0.86 |
| *SLUG* | 10.5 | 7.4 | 1.7 | 2.9 |
| *ZEB1* | 5.3 | 3.4 | 10.1 | 2 |
| *ZEB2* | 17.3 | 1.7 | 8.6 | 2.2 |
| *TWIST1* | 15.2 | 1.4 | 1.4 | 0.76 |
| *TGF-β1* | 10.3 | 1.8 | 11.1 | 2.8 |
| *TGF-β2* | 1.3 | 0.5 | 10.9 | 18.67 |
| *PAR3* | 0.18 | 0.82 | 1.1 | 0.28 |
| *PALS* | 0.06 | 0.56 | 0.09 | 0.06 |
| *SCRIB* | 0.47 | 0.68 | 0.01 | 0.06 |
| **Relative expression levels of select genes determined by immunostaining** |  |  |  |  |
| KRT12 | DR | DR | DR | NT |
| E-Cadherin | DR | UR | UR | NT |
| β-Catenin (membrane) | DR | DR | DR | NT |
| β-Catenin (nuclear) | UR | UR | UR | NT |
| Survivin | UR | UR | UR | NT |
| Ki67 | UR | UR | UR | NT |
| PAR3 | DR | DR | DR | NT |
| PALS1 | DR | DR | DR | NT |
| SCRIB | DR | DR | DR | NT |
| KLF4 | DR | DR | DR | NT |

**Supplementary Table S2.**
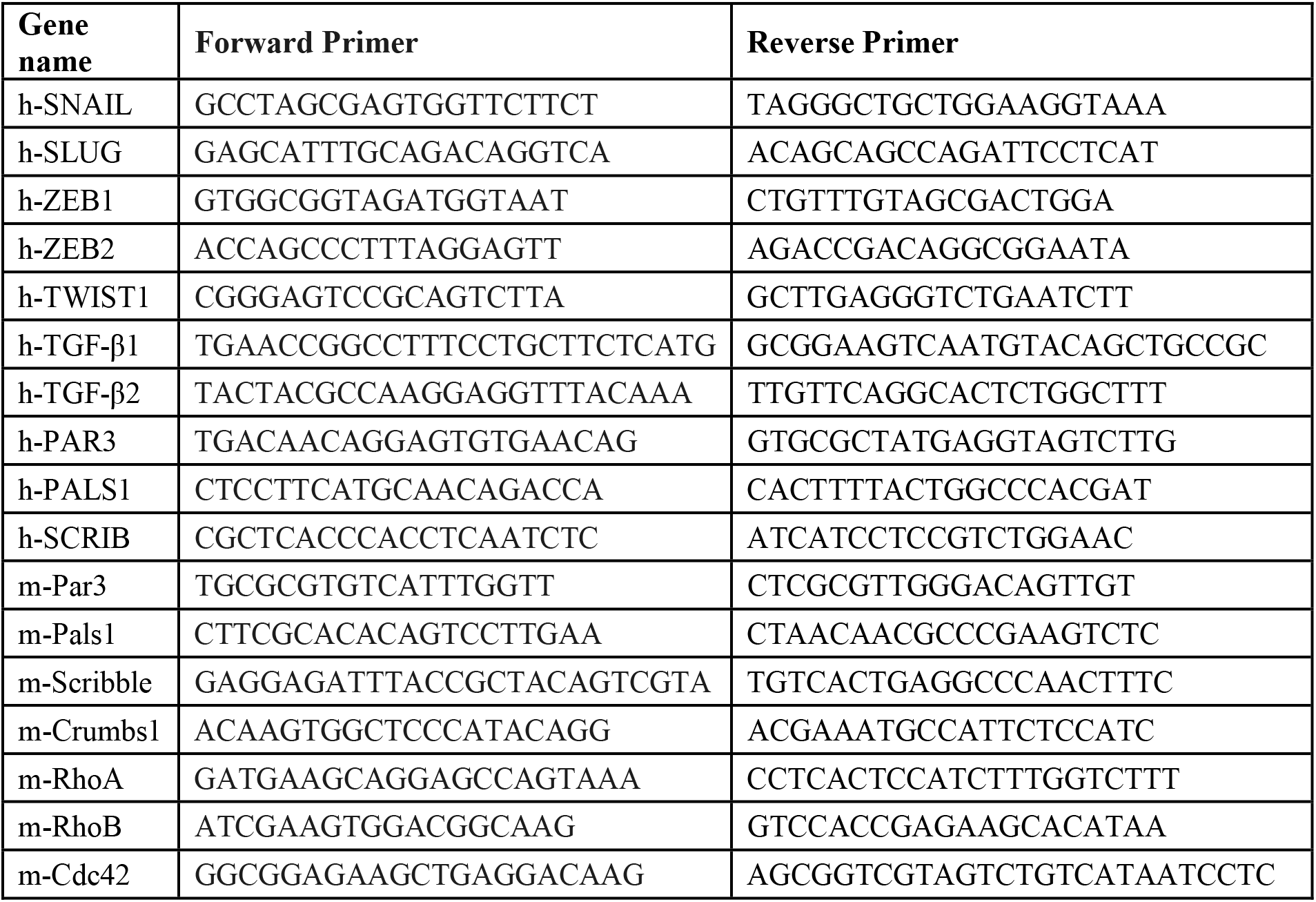
Sequence of oligonucleotide primers used. Prefix h indicates human genes and m, mouse genes.

**Supplementary Table S3.**
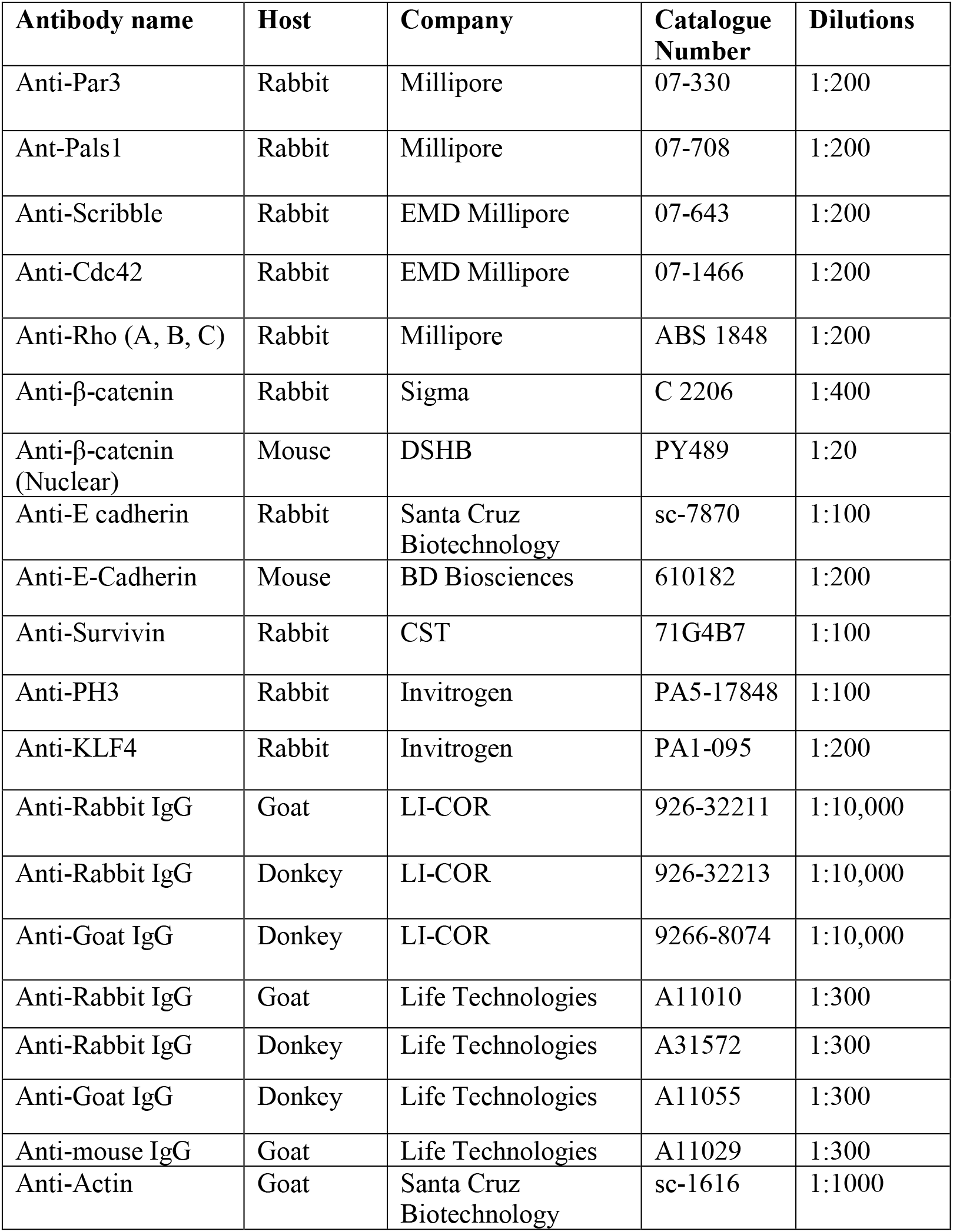
List of antibodies used.

| **Antibody name** | **Host** | **Company** | **Catalogue Number** | **Dilutions** |
| --- | --- | --- | --- | --- |
| Anti-Par3 | Rabbit | Millipore | 07-330 | 1:200 |
| Ant-Pals1 | Rabbit | Millipore | 07-708 | 1:200 |
| Anti-Scribble | Rabbit | EMD Millipore | 07-643 | 1:200 |
| Anti-Cdc42 | Rabbit | EMD Millipore | 07-1466 | 1:200 |
| Anti-Rho (A, B, C) | Rabbit | Millipore | ABS 1848 | 1:200 |
| Anti-β-catenin | Rabbit | Sigma | C 2206 | 1:400 |
| Anti-β-catenin (Nuclear) | Mouse | DSHB | PY489 | 1:20 |
| Anti-E cadherin | Rabbit | Santa Cruz Biotechnology | sc-7870 | 1:100 |
| Anti-E-Cadherin | Mouse | BD Biosciences | 610182 | 1:200 |
| Anti-Survivin | Rabbit | CST | 71G4B7 | 1:100 |
| Anti-PH3 | Rabbit | Invitrogen | PA5-17848 | 1:100 |
| Anti-KLF4 | Rabbit | Invitrogen | PA1-095 | 1:200 |
| Anti-Rabbit IgG | Goat | LI-COR | 926-32211 | 1:10,000 |
| Anti-Rabbit IgG | Donkey | LI-COR | 926-32213 | 1:10,000 |
| Anti-Goat IgG | Donkey | LI-COR | 9266-8074 | 1:10,000 |
| Anti-Rabbit IgG | Goat | Life Technologies | A11010 | 1:300 |
| Anti-Rabbit IgG | Donkey | Life Technologies | A31572 | 1:300 |
| Anti-Goat IgG | Donkey | Life Technologies | A11055 | 1:300 |
| Anti-mouse IgG | Goat | Life Technologies | A11029 | 1:300 |
| Anti-Actin | Goat | Santa Cruz Biotechnology | sc-1616 | 1:1000 |

